# Genome Diversity in Ukraine

**DOI:** 10.1101/2020.08.07.238329

**Authors:** Taras K. Oleksyk, Walter W. Wolfsberger, Alexandra Weber, Khrystyna Shchubelka, Olga T. Oleksyk, Olga Levchuk, Alla Patrus, Nelya Lazar, Stephanie O. Castro-Marquez, Patricia Boldyzhar, Alina Urbanovych, Viktoriya Stakhovska, Kateryna Malyar, Svitlana Chervyakova, Olena Podoroha, Natalia Kovalchuk, Yaroslava Hasynets, Juan L. Rodriguez-Flores, Sarah Medley, Fabia Battistuzzi, Ryan Liu, Yong Hou, Siru Chen, Huanming Yang, Meredith Yeager, Michael Dean, Ryan E. Mills, Volodymyr Smolanka

**Author notes:** Corresponding address: Dr. Taras K. Oleksyk. or Dr. Ryan Mills. these authors contributed equally (Walter W. Wolfsberger, Alexandra Weber, Khrystyna Shchubelka).

## Abstract

The main goal of this collaborative effort is to provide genome wide data for the previously underrepresented population in Eastern Europe, and to provide cross-validation of the data from genome sequences and genotypes of the same individuals acquired by different technologies. We collected 97 genome-grade DNA samples from consented individuals representing major regions of Ukraine that were consented for the public data release. DNBSEQ-G50 sequences, and genotypes by an Illumina GWAS chip were cross-validated on multiple samples, and additionally referenced to one sample that has been resequenced by Illumina NovaSeq6000 S4 at high coverage. The genome data has been searched for genomic variation represented in this population, and a number of variants have been reported: large structural variants, indels, CNVs, SNPs and microsatellites. This study provides the largest to-date survey of genetic variation in Ukraine, creating a public reference resource aiming to provide data for historic and medical research in a large understudied population. While most of the common variation is shared with other European populations, this survey of population variation contributes a number of novel SNPs and structural variants that have not been reported in the gnomAD/1KG databases representing global distribution of genomic variation. These endemic variants will become a valuable resource for designing future population and clinical studies, help address questions about ancestry and admixture, and will fill a missing place in the puzzle characterizing human population diversity in Eastern Europe. Our results indicate that genetic diversity of the Ukrainian population is uniquely shaped by the evolutionary and demographic forces, and cannot be ignored in the future genetic and biomedical studies. This data will contribute a wealth of new information bringing forth different risk and/or protective alleles. The newly discovered low frequency and local variants can be added to the current genotyping arrays for genome wide association studies, clinical trials, and in genome assessment of proliferating cancer cells.

## Data Description

### The context

Ukraine is the largest country located fully in Europe with a population that was formed as a result of several millennia of migration, and admixture. It occupies the intersection between the westernmost reach of the great steppe and the easternmost extent of the great forests that spread across Europe, at the crossroad of the great trade routes from “Variangians to the Greeks’’ along the river Dnipro, which the ancient Greeks referred to as Borysthenes, and the Silk Road linking civilizations of Europe and Asia [1]. This land has seen the great human migrations of the Middle Ages sweeping from across the great plains, and even before that, in the more distant past, the early farmers [2] and the nomads who first domesticated the horse [3–6]. Here, at the dawn of modern human expansion, our ancestors met the Neanderthals who used to hunt the great game along the glacier of the Ice Age [7,8].

The rich history shaped genetic diversity among the people living in the country today. As people have moved and settled across this land, they have contributed unique genetic variation that varies across the country. While the ethnic Ukrainians constitute approximately more than three quarters of the total population of modern Ukraine, this majority is not uniform. A large Russian minority compose approximately one-fifth of the total population with higher concentration in the southeast. Smaller minority groups are historically present in different parts of the country: Belarusians, Moldovans, Bulgarians, Poles, Jews, Greeks, Hungarians, Romanians, Roma (Gypsies), and others [9].

This study offers genome data from 97 individuals from Ukraine (Ukrainians from Ukraine or UAU) to the scientific community in order to help fill the gaps in the current knowledge about the genomic variation in Eastern Europe, a part of the world that has been largely and consistently overlooked in the global genomic surveys [10]. This was the first effort to describe and evaluate the genome wide diversity in Ukraine. Samples were successfully sequenced using BGI’s DNBSEQ™ technology, and cross-validated by Illumina sequencing and genotyping. The major objectives of this study was to demonstrate the importance of studying local variation in the region, and to demonstrate the distinct and un que genetic components of this population. Of particular interest were the medically related variants, especially those with allele frequencies that differed with the neighboring populations. As a result, we present and describe an annotated dataset of genome-wide variation in genomes of healthy adults sampled across the country.

### The dataset

The new dataset includes 97 whole genomes of self-reported Ukrainians from Ukraine at 30x coverage sequenced using DNBSEQ-G50 (formerly known as BGISEQ-500; BGI Inc., Shenzhen, China) and annotated for genomic variants: SNPs, indels, structural variants and mobile elements. The samples have been collected across the entire territory of Ukraine, after obtaining the IRB approval (Protocol #1 from 09/18/2018, **Supplementary File 1**) for the entire study design, and informed consent from each participating volunteer (**Supplementary File 2**). Each participant in this study had an opportunity to review the informed consent, have been explained the nature of the genome data, and made a personal decision about making it public.

The majority of samples in this study (86 out of 97) were additionally genotyped using Illumina Global Screening Array (Illumina Inc., San Diego, USA) in order to confirm the accuracy of base calling between the two platforms. In addition, one sample (EG600036) was also sequenced on the Illumina HiSeq (∼60x coverage) and also used for validation of the variant calls (see summary in **Table S1**, and full sequencing statistics for individual samples in **Table S1**.**2**). The list of the cross validated samples and the source technology of the data is presented in the **Supplementary File 3**.

The current dataset contains locations and frequencies of more than 13M unique variants in Ukrainians from Ukraine (**UAU**) which are further interrogated for functional impact and relevance to the medically related phenotypes (**Table 1, Supplementary Data 4**). As much as 3.7% of these alleles, or 478 K, are novel genomic SNPs that have never been previously registered in the gnomAD database [11] (**Table 1**). This number is similar in magnitude to what was reported earlier in two populations from European Russia (3-4%; [12]). Many of the discovered variants (12.6%) are also currently missing from the global survey of genomic diversity in the 1,000 Genomes Project [13]. Majority of these described variants are rare or very rare (<5%; **Figure S2**).

**Table 1.**
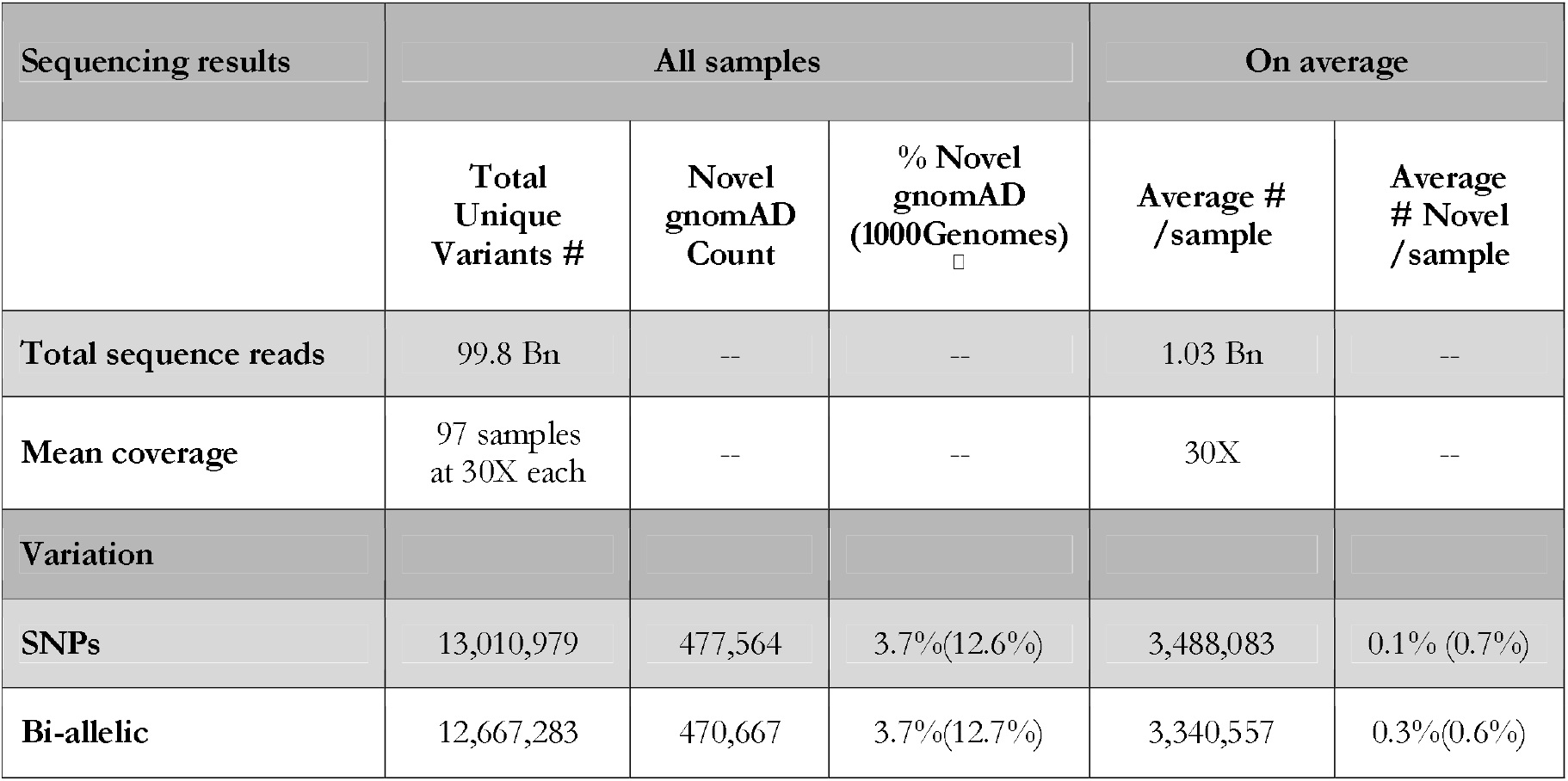
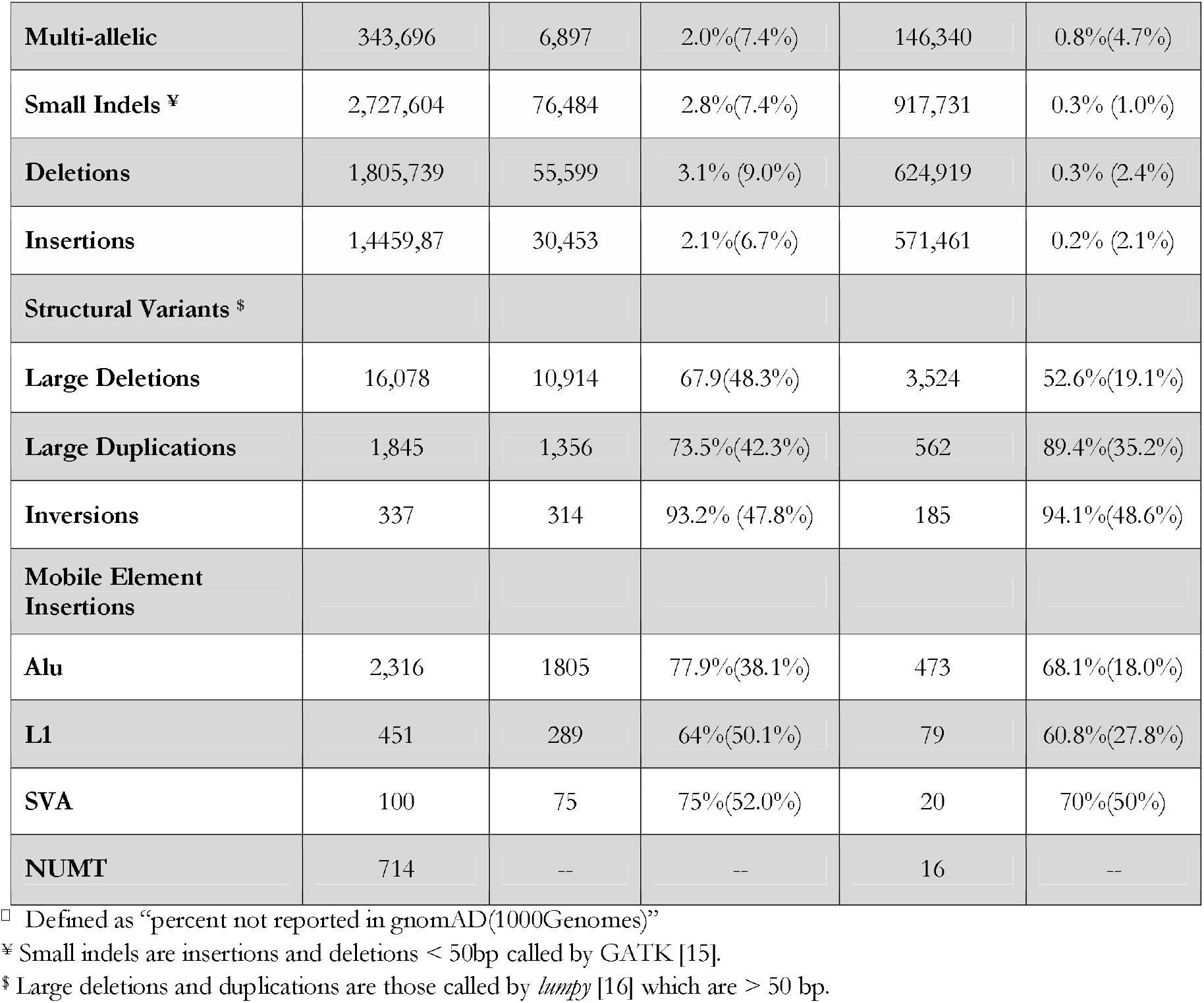
Summary of variation in the 97 whole genome sequences from Ukraine.

Unless other indigenous ethnic groups from Ukraine (such as the Crimean Tatars), would be included in the study, increasing the sample size above from 100 to 1,000 individuals is not likely to greatly contribute to discovery of novel mutations [14]. The proportion of the novel structural variants and mobile elements compared to the earlier databases is even higher: almost 1M (909,991) complex indels, regions of simultaneous deletions and insertions of DNA fragments of different sizes which lead to net a change in length, majori y of which are novel (Table 1). Many of the newly discovered variants are functional and potentially contribute to the phenotype (classified in Table 2). We report many important variants that are overlooked or require special modifications in the commonly used resources and tools in genomic research and diagnostics. This wealth of novel variation underscores the importance of variant discovery in local populations that cannot be ignored in biomedical studies.

**Table 2.**
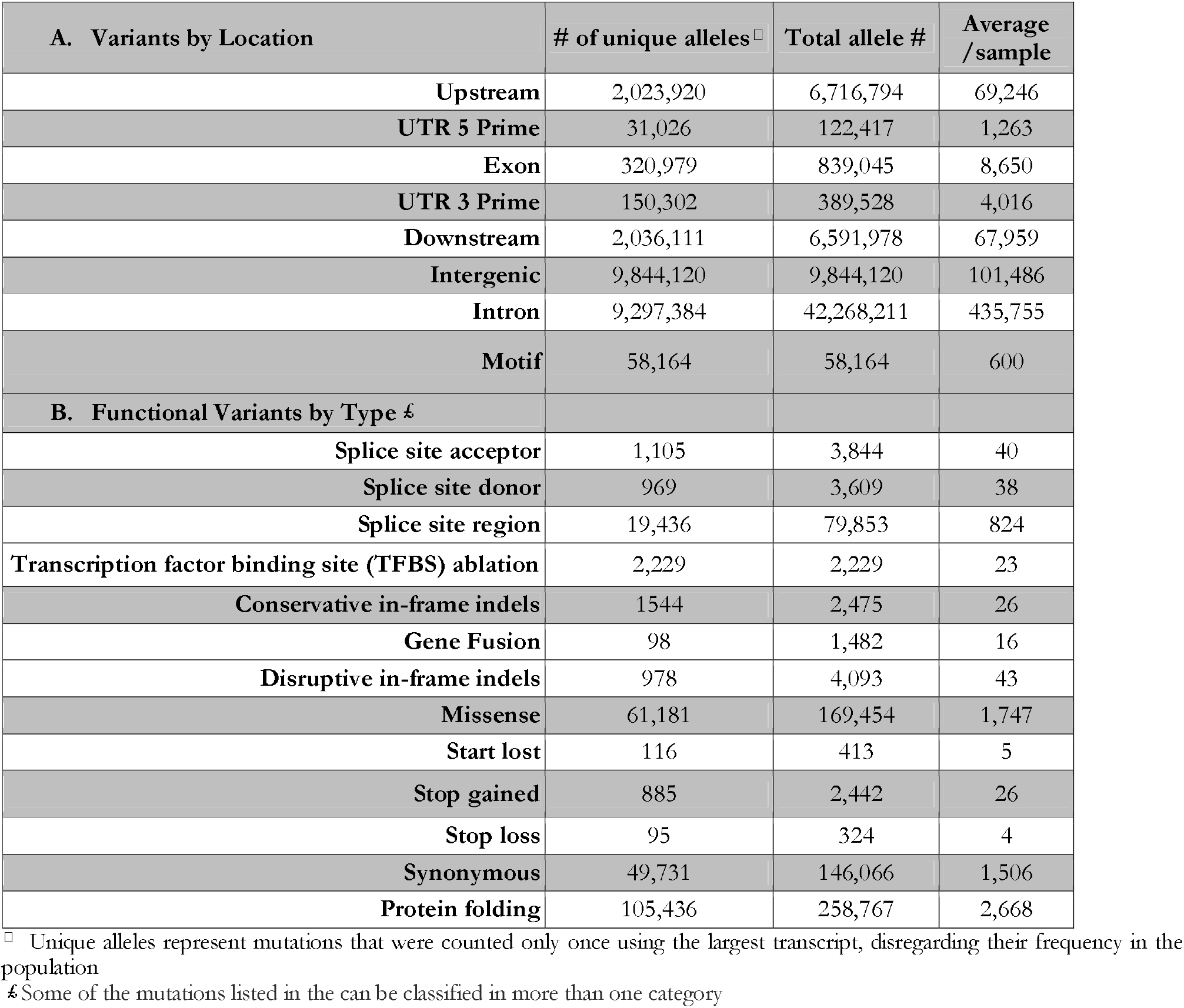
Summary annotation of different genomic elements in the Ukrainian genomes annotated in BGISeq data from 97 Ukrainian samples

### Variant calling and confirmation

For each sample in the database, we estimated the number of passing bi-allelic SNPs calls (i.e. loci with the non-reference genotypes relative to the most current major human genome assembly, GRCh38 [17])(**Table 1**). Approximately 12% of these were filtered out based on excess heterozygosity and low variant quality scores (**Table S2**). For the indels, we also estimated the number of passing calls compared to GRCh38, and excluded 4% of those which did not pass filtering. The total number of the unique SNPs, small and large indels (**Table 1**) was calculated from the raw reads alignments of all the 97 sequenced genomes (**Total Unique SNPs, Table S2**) with the exception of those filtered out for low variant quality scores and containing excess heterozygosity (**Filtered Count; Table S2**). In addition, we filtered out 4,135,903 variants that only appeared once in a single sample (for both indels and SNPs) and designated them as “*singletons*”.

We report a good correspondence between the SNP calls made using DNBSEQ and NovaSeq data. A comparison of the variants detected using these three platforms for sample EG600036 are summarized in **Figure 1.A**. The SNP concordance for samples with both DNBSEQ and SNP array data is summarized in **Figure 1C**. The cross-platform comparison shows a very good overlap across all three technologies: with more than 3.5 M SNPs (or **97**.**7**%) of the SNPs identified in the DNBSEQ were also verified in the whole genome sequence of EG600036 sequenced by the Illumina NovaSeq. The correspondence with the Illumina SNP Array for sample EG600036 was also very good: **95**.**8**% of all the SNPs genotypes called by the Illumina method were also detected by the DNBSEQ (**Figure 1.A(Right), C(Right)**). The concordance between the non-reference alleles between the two platforms in all the 86 samples was nearly linear (*r2* =0.985, **Figure 1.C(Left)**).

**Figure 1.**
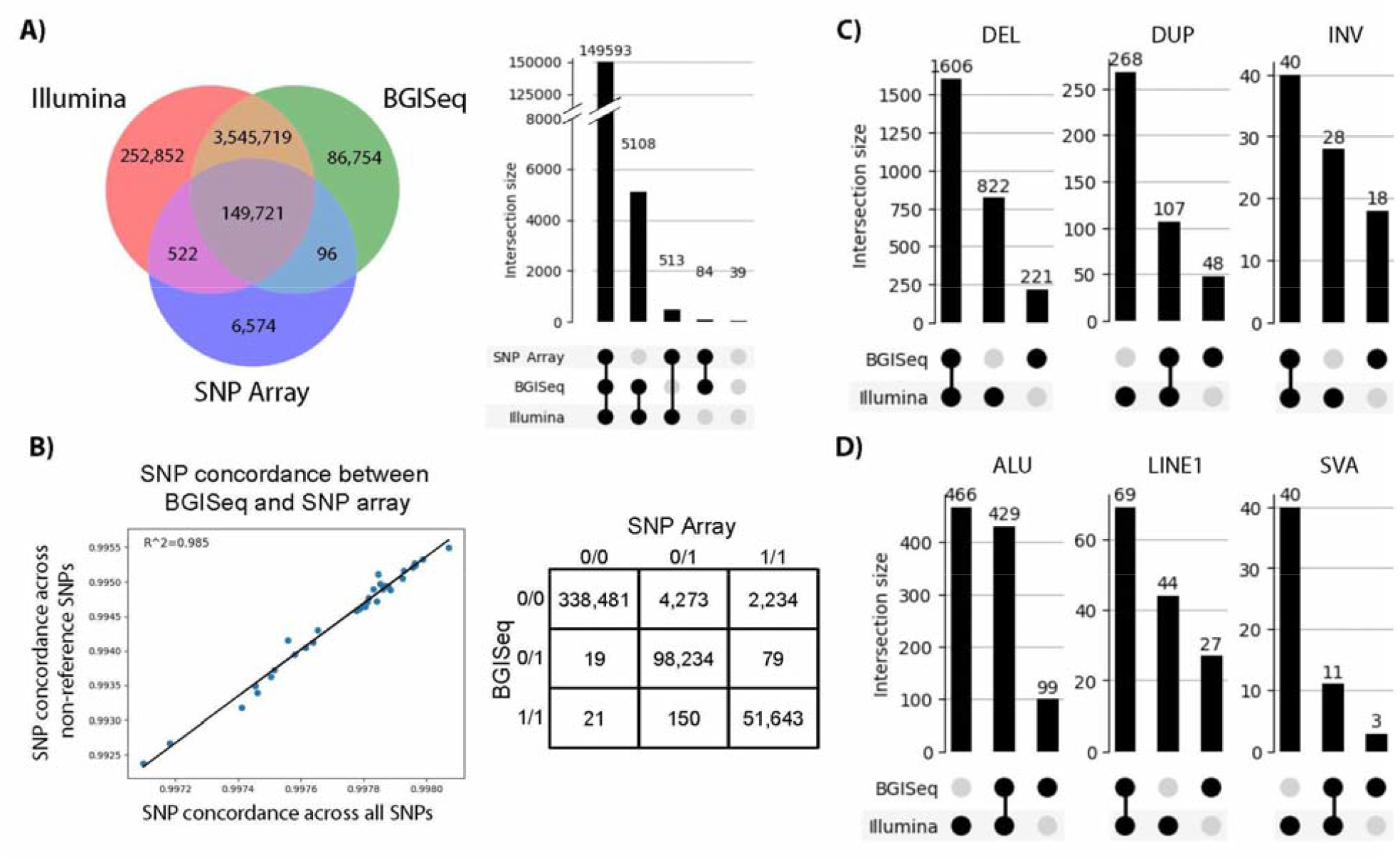
Variant concordance across the three sequencing/genotype methods. **A) Left:** Overlap of SNP positions identified in one sample (EG600036) using each of the three platforms. **Right:** Concordance of SNP genotypes in one sample derived from each of the three platforms. This only includes the subset of SNPs with alternate alleles included in the Illumina genotyping array (the smallest of the three variant sets). The variants indicated as belonging to none of the categories are variants whose genotypes differ between all three platforms. **B) Left:** The percentage (%) of concordance between the Illumina SNP array and DNBSEQ for all SNPs compared to the % concordance of only SNPs with non-reference alleles in the Illumina SNP array for the 86 samples genotyped on both platforms. **Right:** Concordance of SNP genotypes between DNBSEQ and Illumina SNP Array for one sample (EG600036). **C)** Overlap within the numbers of the three major structural variants detected in one sample using the two whole genome sequencing datasets. **D)** Overlap within the numbers of the three major mobile element insertions detected in one sample using the two whole genome sequencing datasets.

Transition/Transversion ratio (or TITV ratio) for the novel SNPs (estimated with *TiTvtools* [18] and visua ized by *plotTiTv* in **Figure S1**) was lower than the TITV ratio for SNPs in the dbSNPs database (**1**.**9 vs 2**.**2**; [19]). Similarly, insertions to deletions (ins/del) ratio for novel indels is lower than for the indels already reported in the dbSNP database (**0**.**63 vs 0**.**75**). This observation likely reflects our improved ability to detect small insertions in newer sequencing technologies compared to many platforms which historically submitted variation to dbSNP.

We have defined the multi-allelic SNPs as observations of genomic positions having two or more alternative alleles [20]. These are important variants that are overlooked or require special modifications in the commonly used resources and tools in genomic research and diagnostics. We report a total of 343,696 multiallelic sites in the sequences from our sample of which 2.0% are at locations unreported in the gnomAD database [11] (**Table 1**).

In addition to the SNPs, we have identified and quantified major classes of structural variations in the Ukrainian population: small indels (insertions and deletions < 50bp), large structural variants (deletions, duplications and inversions > 50 bp) and Mobile Element Insertions (MEI)(Alu-s, L1 elements, non-autonomous retroelements (SVA), and nuclear mitochondrial DNA (NUMT) copies). A number of structural elements were reported, including common and novel ones. While among the small variants most were common (6-9%), a large proportion of large variants and MEIs (38-52%) have not been reported previously in the 1000Genomes Database (**Table 1**).

Once more, there is a significant correspondence between the calls made using BGI DNBSEQ and Illumina NovaSeq data. The two sequencing platforms show a significant overlap in calling indels (**DEL**): 87.9% of the variants called by the DNBSEQ were also detected by the Illumina. At the same time, there were 822 deletions, or 33.8% of all the indels called by the Illumina that were not detected by the DNBSEQ (**Figure 1.B**). A similar picture, where DNBSEQ performs competitively well, is also observed for inversions (**INV**)(**Figure 1.B**), and **LINE1** transposable elements (**Figure 1.D**). At the same time, more Duplications (**DUP**)(**Figure 1.B**), and the two classes of transposable elements evaluated: Alu elements (**ALU**) and the non-autonomous retroelements (**SVA**)(**Figure 1.D**). Evaluation tests show that current algorithms are platform dependent, in the sense that they exhibit their best performance for specific types of structural variation as well as for specific size ranges [21], and the algorithms designed for detection and archived datasets are predominantly for Illumina pair-end sequencing [22,23]. While it is possible that these results indicate Illumina’s superiority at detecting structural variation, it also can also be the consequence of the bioinformatics tools for calling structural variants developed using mainly the Illumina data, as suggested by previous comparative evaluations of the two technologies [24,25]. Additionally, higher coverage of the Illumina data (60x) could have contributed to the differences observed between the platforms.

The database was compared to the existing global resources of population variation such as Genome Aggregation Database (gnomAD)[11] and the 1000Genomes Project (1KG)[13]. Specifically, under our search criteria, the small variants (SNPs and Small Indels) were considered “*novel*” if they were absent from all the samples in the two global datasets (gnomAD and 1KG; **Table 1**). The large structural variants and Mobile Element Insertions were considered “novel” if the variant was not present in the gnomAD and 1KG databases. To determine if a given variant was present in one of the databases, a variant of the same type in the database had to overlap the Ukrainian variant with a minimum fraction of 0.95. We observed no significant deviation of the rate at which reference bases were observed at REF/alt heterozygous SNP sites (reference bias was near 50%).

### Collection of functional variants

A particular interest in this study is the distribution of functional variation, not in the least due to the potential impact on phenotypes, especially to those with medical relevance [26]. As much as 97.5% of all annotated variation was discovered outside of the known functional elements (upstream, downstream, intron and intergenic). These results are similar to the expected distributions of mutations shown with the simulated data [27]. Nevertheless, there were more than 8,000 mutations discovered within exons of each individual on average (**Table 2**.**A**). We annotated several classes of functional mutations within the coding regions (**Table 2**.**B**). As expected, the nonsense mutations classified in the annotation file as “*Disruptive in-frame indel*”, “*Start lost*”, “*Stop gained*”, and “*Stop loss*” were rare, while categories with minimal effect on the function, such as “*Synonymous*”, “*Motiff*”, “*Protein folding*”, “*Missense*” were more common. Some of the mutations listed in the can be classified in more than one category (e.g. “*Synonymous variants*” can also be counted in “*Exonic variants*”).

In addition to the gene coding mutations, we report a number of regulatory variants. For example, the database contains a total of 2,229 transcription factor binding site ablation (TFBS) mutations (**Table 2**.**B**). A summary of functional variation discovered in this study is presented in **Table 2**. The full list of high impact functional variants (including frameshift, start lost/stop lost or gained, transcript ablations and splice alterations) that had an allele count of two or more with their predicted function, number of gene transcripts of the gene affected, and frequencies is presented in **Table S3**. The full annotation database with classifications is available online as **Supplementary File 4** *(GigaScience ftp://user81@8.210.79.81/Ukraine_bgi_all_ann_GWAS_Clinvar.vcf.gz)*

### Collection of the medically relevant variants

Many of the reported variants are already known to be medically related, and are listed either in Genome-wide association studies (GWAS) [28] or ClinVar (a NCBI archive of reports of the relationships among human variations and phenotypes with supporting evidence) [29] catalogues (**Table 3**). Our database contains a total of 43,892 benign mutations in medically related genes, but also 189 unique pathogenic or likely pathogenic variants, as well as 20 protective or likely protective alleles as defined in ClinVar [29,30]. Each individual in this study carries 19 pathogenic and 12 protective mutations on average. While least some individuals were homozygous for the pathogenic allele, none of the associated disease phenotypes have been reported, which could be largely attributed to heterozygosity, age-dependent penetrance, expressivity and gene-by-environment interactions [31,32].

**Table 3.** Medically-relevant variants in the Ukrainian population included in GWAS [28] and ClinVar [29] databases

| Source of Annotation | # Unique substitutions <sup>□</sup> | Total allele # | Average /sample |
| --- | --- | --- | --- |
| **GWAS catalog** | 102551 | 6,479,953 | 66804 |
| **ClinVar: pathogenic (or likely pathogenic)** | 189 | 1,830 | 19 |
| **ClinVar: benign (or likely benign)** | 43,892 | 1,842,668 | 18997 |
| **ClinVar: protective (or likely protective)** | 20 | 1,209 | 12 |

<sup>□</sup> Unique variants represent substitutions that were counted only once, disregarding their frequency in the population

As expected, our study shared a lot more variants with the GWAS [28] than with the ClinVar [29] catalogue. While GWAS has recently become the tool of choice to identify genetic variants associated with complex disease and other phenotypes of interest [33], since the amount of genetic variance explained by these variants is low, they are generally not very useful for prediction pathogenic phenotypes [34]. It is also important to note, that not all ClinVar variants carry the same weight of supporting evidence, attributing disease causation to prioritized variants remains an inexact process and some of the reported associations eventually are proven to be spurious [35]. Nevertheless, the importance of the unique set of mutations published here is difficult to overemphasize, as it constitutes the first published set of pathological variants in an understudied population, an important step towards a local catalogue of medically relevant mutations. In addition, as the attention in the genomic community is shifting from monogenic to polygenic traits, many of these may become relevant in the future research and exploration [36]. Full list of the medically relevant functional markers found in the Ukrainian population and reported in GWAS [28] and ClinVar [29] databases. with alternative allele frequencies and annotations are presented in **Tables S4**.

Disease variants with frequencies that differed between the Ukrainians and the neighboring populations are of particular interest to the medical community. It is well established that differences in allele frequencies are a consequence of evolutionary forces acting in populations (such as drift, mutation, migration, nonrandom mating and natural selection), the certain diseases and heritable traits display marked differences in frequency between populations [37]. With this in mind, we created a list of the known disease variants that whose frequencies differ between Ukrainians and other European populations (the combined European sample (EUR) from the 1000Genomes Project (Utah Residents (CEPH) with Northern and Western European Ancestry, Toscani in Italy (TSI), Finnish in Finland (FIN) British in England and Scotland (GBR), Iberian Population in Spain (IBS)[13,38] and French population from HGDP (FRA)[39]) and Russians from HGDP (RUS)[39]. Several examples of these variants are presented in **Table 4**. Among these are variants involved in a number of medical conditions such as hyperglycinuria/iminoglycinuria (rs35329108; SLC6A19), efficacy of bisphosphonate response (rs2297480; FDPS), autism (rs7794745, CNTNAP2), Leber congenital amaurosis (rs10151259, RPGRIP1), and breast cancer susceptibility in BRCA1 and BRCA2 carriers n (rs1801320, RAD51)(**Table 4**). Of course, not all the medically related variants are currently known, and many remain to be discovered and verified in local populations. This is, to some extent, a consequence underreporting of allelic endemism w thin understudied populations, particularly in Eastern Europe [10] but also elsewhere [40,41]. By offering public annotations of functional mutations in a population sampled across the territory of Ukraine, our database contributes a number of candidates to direct future research in medical genomics. We chose only the **markers with the highest non-reference allele frequency (NAF) differences** compared to the neighboring populations: the combined population from Europe (EUR; [13]) and Russians from HGDP (RUS)[39] evaluated by the Fisher Exact Test (FET) and listed them **Table 5**.

**Table 4.** Examples of the functional SNPs with highly differentiating functional markers reported in ClinVar [29], with high differences in the Ukrainian population compared to the neighboring populations in other European populations (the combined sample from Western and Central Europe from 1000Genomes Project (EUR)[13,38] and French population from HGDP (FRA)[39], as well as Russians (RUS) from HGDP [39]. Non-reference allele frequency (NAF) is reported compared to the reference allele in GRCh38. Differences are evaluated by the Fisher Exact Test (FET). All the functional SNPs with significant population frequency differences are listed in **<u>Table S5</u>**.

| SNP | Chr | Gene | REF/alt<sup>□</sup> | Associated medical condition | NAF UKR | NAF EUR | NAF RUS | FET vs. EUR (p-value) | FET vs. RUS (p-value) |
| --- | --- | --- | --- | --- | --- | --- | --- | --- | --- |
| *rs2297480* | 1 | *FDPS* | T/G | Efficacy of the Bisphosphonate response | 0.13 | 0.23 | 0.27 | 0.038 | >0.001 |
| *rs35329108* | 5 | *SLC6A19* | G/A | Hyperglycinuria. Iminoglycinuria | 0.32 | 0.26 | 0.17 | 0.049 | 0.004 |
| *rs7794745* | 7 | *CNTNAP2* | A/T | Autism | 0.48 | 0.22 | 0.30 | 0.032 | 0.010 |
| *rs10151259* | 14 | *RPGRIP1* | G/T | Leber congenital amaurosis<br>Cone-rod. Dystrophy | 0.32 | 0.66 | 0.11 | 0.003 | 0.014 |
| *rs1801320* | 15 | *RAD51* | G/C | Breast cancer susceptibility in *BRCA1* and *BRCA2* carriers | 0.19 | 0.31 | 0.07 | 0.047 | 0.000 |

<sup>□</sup> The reference allele is set according to the reference allele in GrCH38.p13 [17].

**Table 5.** Examples of the functional markers with the highest **non-reference allele frequency (NAF) differences** in the Ukrainian population evaluated by the Fisher Exact Test (**FET**) compared to the frequencies in the neighboring populations: the combined population from Europe (**EUR;** [13]) and Russians from HGDP (RUS) [39].

| SNP | Chr | Gene | Ref/Alt | Function | NAF UKR | NAF EUR | NAF RUS | FET vs. CEU (p-value) | FET vs. RUS (p-value) |
| --- | --- | --- | --- | --- | --- | --- | --- | --- | --- |
| *rs72625995* | 17 | *POM121L8P* | *C/T* | *exonic , nonsynonymous SNV* | *0.03* | *0.62* | *0.75* | *2.50E-07* | *1.86E-06* |
| *rs9930886* | 16 | *PTPRN2* | *A/G* | *exonic, synonymous SNV* | *0.01* | *0.33* | *0.35* | *2.56E-07* | *2.19E-06* |
| *rs4779816* | 15 | *ZBTB9; BAK1* | *A/G* | *exonic , nonsynonymous SNV* | *0.41* | *0.80* | *0.83* | *3.29E-06* | *7.82E-07* |
| *rs58580222* | 12 | *ABCC1* | *G/A* | *exonic , synonymous SNV* | *0.03* | *0.13* | *0.26* | *3.06E-04* | *1.17E-02* |
| *rs80150964* | 11 | *SMIM40; KIFC1* | *T/C* | *exonic , non-synonymous SNV* | *0.03* | *0.23* | *0.19* | *4.95E-04* | *1.96E-06* |

### Population structure and ancestry informative markers

We performed several population analyses, but only to demonstrate the uniqueness and usefulness of this new dataset. Our results indicate that genetic diversity of the Ukrainian population is uniquely shaped by the evolutionary and demographic forces and cannot be ignored in the future genetic studies. However, we do not evaluate any historical hypotheses on the timing of origins, founding, migration, and admixture of this population, and use only the naive approaches, choosing models based on the statistical models.

To demonstrate the extent to which our dataset contributes to the genetic map of Europe, we explored genetic relationships between Ukrainian individuals within our sample and evaluated genetic differences between this population and its immediate neighbors on the European continent for which population da a of full genome sequences was publicly available. A Principal Component Analysis (PCA) of the merged dataset of 654 samples included European populations from the 1000Genomes Project (Utah Residents (CEU) with Northern and Western European Ancestry, Toscani in Italy (TSI), Finnish in Finland (FIN) British in England and Scotland (GBR), Iberian Population in Spain (IBS)) [13,38]), and French and Russians (RUS) populations from the HGDP [39] as well as the relevant high-coverage human genomes from the Estonian Biocentre Human Genome Diversity Panel (EGDP: Croatians (CRO), Estonians (EST), Germans (GER), Moldovans (MOL), Polish (POL), and Ukrainians (UKR)[42], and Simmons Genome Diversity project (Czechs (CZ), Estonians (EST), French (FRA), Greeks (GRE), and Polish (POL) [43] (**Figure 2**). The latter paper also identifies “Cossacks” as a separate self-identified ethnic group within Russians (Cossacks (RUS) or Ukrainians (Cossacks (UKR)) [43] (**Supplementary File 5**).

**Figure 2.**
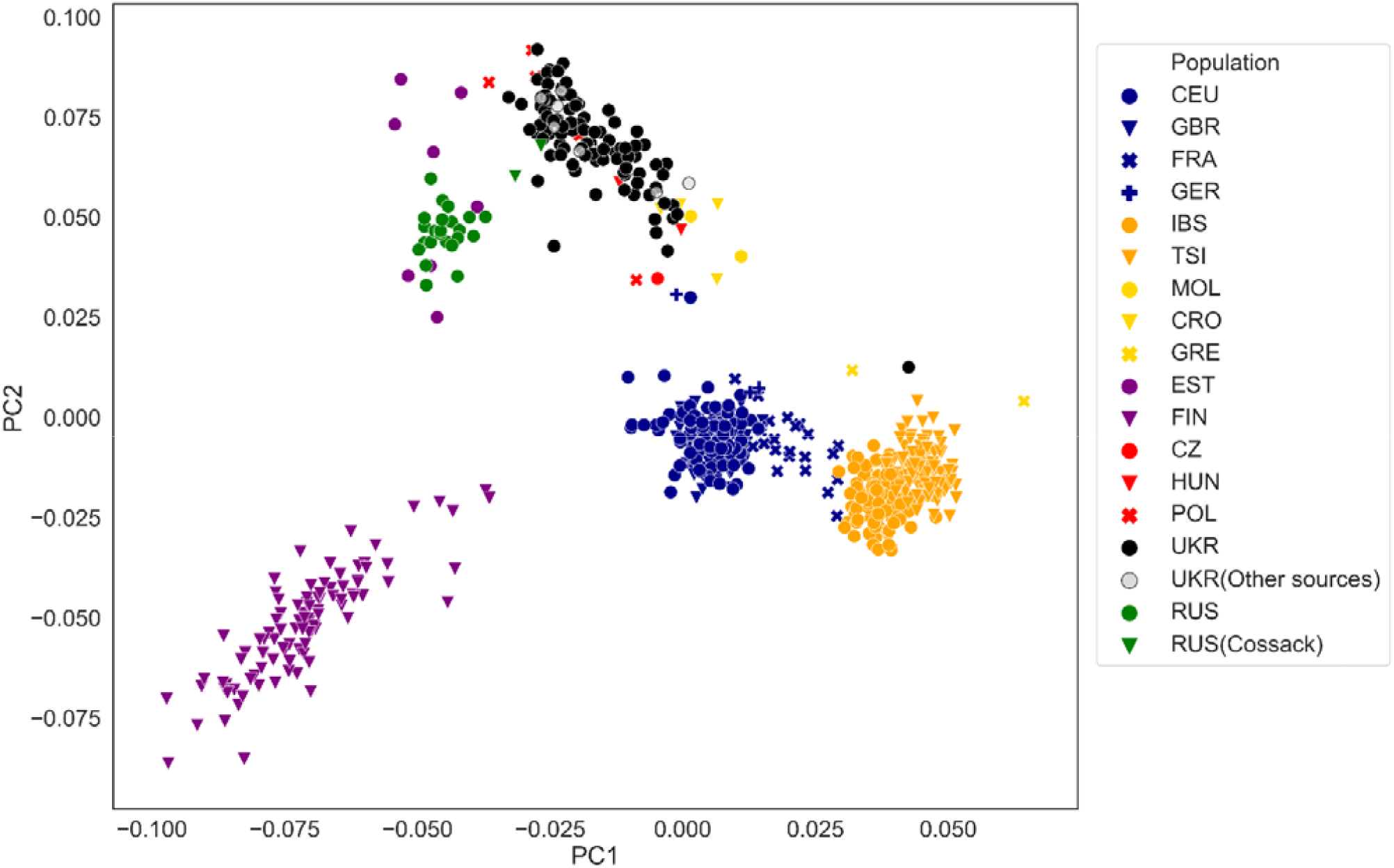
The Principal Component (PC) analysis of genetic merged dataset, containing European populations. Colors reflect prior population assignments from the European samples from the 1000Genomes Project (Utah Residents (CEPH) with Northern and Western European Ancestry, Toscani in Italy (TSI), Finnish in Finland (FIN), British in England and Scotland (GBR), Iberian Population in Spain (IBS)[13,38], French (FRA) and Russians (RUS) from HGDP (RUS) [39] as well as the relevant high-coverage human genomes Croatian (CRO), Czech (CZ), Estonian (EST), German (GER), Greek (GRE), Hungarian (HUN), Moldovan (MOL), Polish (POL), Russian Cossack (RUS) and Ukrainian (UKR) from the Estonian Biocentre Human Genome Diversity Panel (EGDP) [42] as well as Simmons Genome Diversity project [43]. The analysis was performed with *Eigensoft* [47].

Ukrainian genomes from this (**black dots**) as well as other studies (**black circles**) [42,43] form a single cluster positioned between the Northern (Russians (**green circles**), Estonians (**purple circles**) on one side, and Western European populations on the other (**blue shapes are**: CEU, French, British and Germans, **Figure 2)**. There was a significant overlap with the other Central and Eastern European populations, such as Czechs (red dots), and Polish (**red crosses**), and the people from the Balkans (Croats, Greeks and Moldovans; **light orange shapes**). This is not surprising, in addition to the close geographic distance between these populations, this may also reflect the insufficient representation of samples from the surrounding populations (see **Supplementary Data 5**). Similarly, the admixture analysis demonstrates distinctiveness of our dataset, but also demonstrates unique combinations of genetic components that may have shaped this population (**Figure 3** and **Figure S3**).

**Figure 3.**
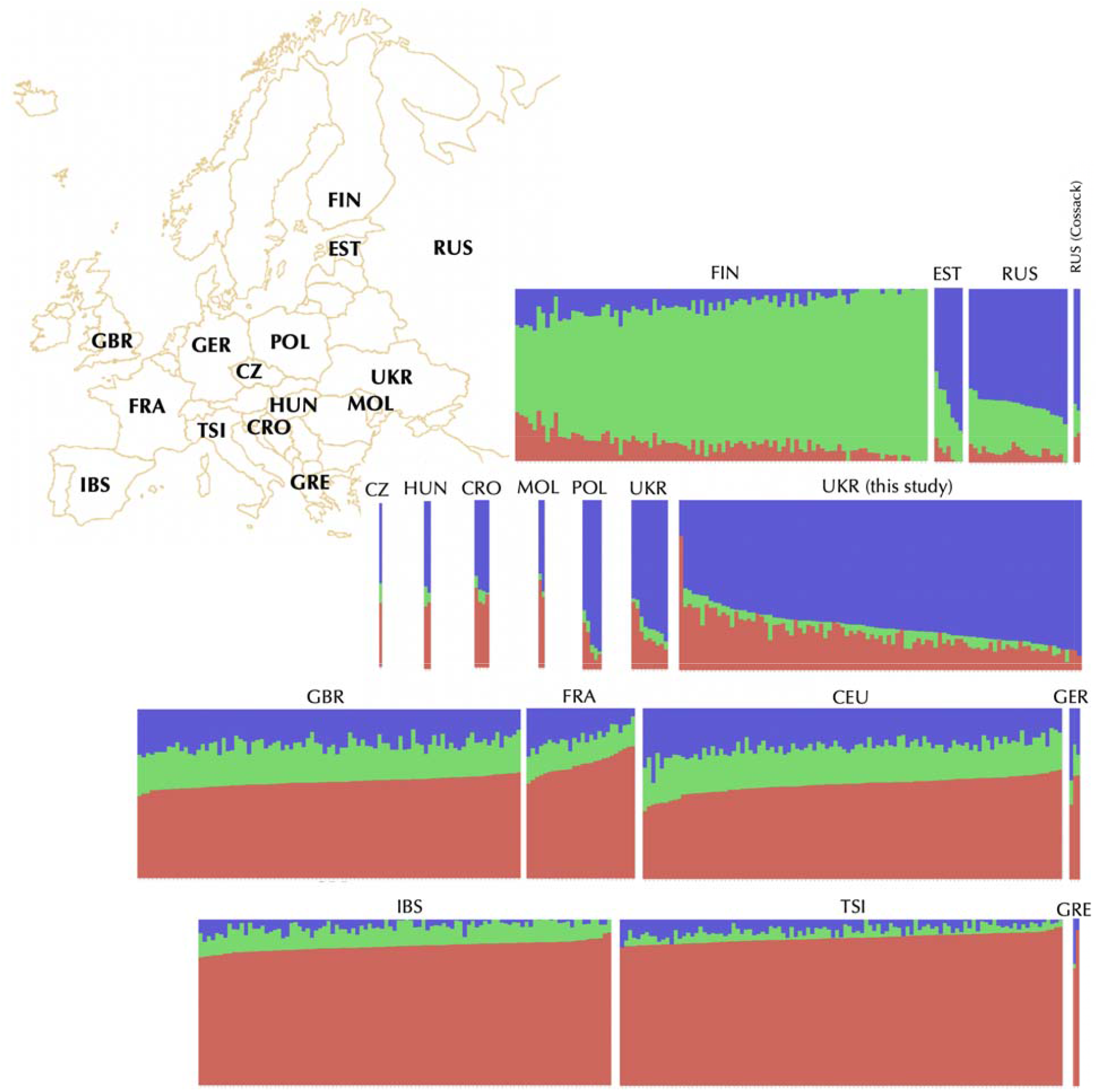
Genetic structure of Ukrainian population in comparison to other European populations. Structure plot constructed ADMIXTURE package [48] at K=3 illustrates similarity and differences between genomes from this study as well as samples from the 1000Genomes Project (Utah Residents (CEU) with Northern and Western European Ancestry, Toscani in Italy (TSI), Finnish in Finland (FIN), British in England and Scotland (GBR), and Iberian Population in Spain (IBS)[13,38], French(FRA) and Russians (RUS) from HGDP [39], as well as the relevant high-coverage human genomes Croatian (CRO), Czech (CZ), Estonian (EST), German (GER), Greek (GRE), Hungarian (HUN), Moldovan (MOL), Polish (POL), Russian Cossack (RUS) and Ukrainian (UKR) from the Estonian Biocentre Human Genome Diversity Panel (EGDP) [42] as well as Simmons Genome Diversity project [43]. For identification of the optimal K parameter, we evaluated a range from 2 to 8, with K=3 resulting in the lowest error. Plots with K=3 to K=6 are presented in **Figure S3**.

Addition of the new genomic data will most likely add to the resolution of the genetic map of this region and further reveal differences between the populations of Eastern and Central Europe. Meanwhile, our dataset showed a limited amount of inbreeding (**Figure S4**) and contains information for future population studies. A list of all the variants with significant difference in frequencies between Ukrainians and other European populations are listed in **Table S6**. This database can be a starting point for association studies, as ancestry informative markers (AIMs)[44], and to be used for mapping disease alleles by admixture disequilibrium [45,46].

To provide a more extended view of the genetic components contributing to the Ukrainian population, we used the population structure plots using the ADMIXTURE package [48]. This allowed us to construct a preliminary picture of putative ancestry contributions and population admixture. In order to identify the optimal K, we implied the 10-fold cross-validation function in range from K=2 to 6. The results with the optimal K=3 shown in **Figure 3** illustrate similarity and the difference of Ukrainian population compared to the other populations in Central and Eastern Europe (**Figure 3, second row**). While the higher values of K (K=3-8; **Figure S3**) show an increasing number of clusters, they also show an increasing amount of error in the cross validation function. This analysis already shows the potential of the current database in helping to resolve population structure in Eastern Europe, but additional genome wide data from neighboring populations would be very helpful to refine the picture in this geographical region. Unfortunately, valuable genome wide data collected from three populations in Russia has been retracted from public databases after the publication [12].

Despite the fact that all of the samples were collected from self-identified ethnic Ukrainians, there were two notable outliers. Sample EG600048 that clustered with the Southern Europeans (Iberia and Italian populations), and EG6000xx clustered with the Western Europeans (CEU, French, British and Germans) (**Figure 2**). This illustrates an important point that while ignoring the unique composition of this population will result in ascertainment bias in biomedical studies. Genetics is not a reliable determinant of ethnicity, but can be used to evaluate individual contributions of ancestry. In anticipating the future ancestry studies we contribute the full list of candidates for Ancestry Informative Markers differentiating Ukrainians with their neighboring populations in Europe (**Table S6**).

People of Ukraine carry many previously known and several novel genetic variants with clinical and functional importance that in many cases show allele frequencies different from neighboring populations in the rest of Europe, including Poland to the West, Romania to the South, the Baltics to the north and Russia to the northeast. While several large genome projects already exists contributing to the understanding of the global genetic variation, many of the rare and endemic alleles that have not been yet identified by the international databases such as the 1,000 Genomes project, and currently not available in standard genotyping panels for association testing for human diseases, and glaring white spots still exists on the genetic maps in local populations of Eastern Europe [10]. We fully expect that the future sampling and sequencing will continue to improve and complete the detailed picture of genomic diversity in people across the country and contribute to the further development of genetic approaches in biomedical research and applications.

## Methods

### a) Sampling strategy

The collection procedure was approved as part of the “*Genome Diversity in Ukraine* ‘’ project by the Institutional Review Board (IRB) of Uzhhorod National University in Uzhhorod, Ukraine (Protocol #1 from 09/18/2018, Supplementary File 1). We employed doctors and medical professionals from different regions of Ukraine to oversee collection of blood samples at hospitals. Healthy (non-hospitalized) volunteers were contacted through advertisements, and invited for personal interviews at outpatient offices. During the visit the volunteers were familiarized with the study and the collection procedure, and gave full consent to participate and have their genotypic and phenotypic data to be freely and publicly available. During each interview, the volunteer participants also completed a questionnaire indicating self-reported region of origin, place of birth of both grandparents (if remembered), sex and several phenotypical features, such as daily history of disease (**Supplementary File 3**). The hard copies of the consents and personal interviews remain sealed and stored at the Biology Department of Uzhhorod National University. After the conclusion of the interview and sample collection, all personal identifiers were removed from the vials containing blood samples, except for an alphanumeric identifier and a barcode. All the subsequent analysis and publication was done in a blind design where neither the participants nor the researchers could identify the person who donated the sample.

At the conclusion of the interview a whole blood sample was collected from a vein into two a 5 ml EDTA tubes by a certified nurse or a phlebotomist, assigned a barcode number, and shipped by courier on dry ice to a biomedical laboratory certified to handle blood samples in Uzhhorod, Ukraine (**<u>Astra Dia Inc</u>.**) for DNA extraction immediately on arrival. The excess of the blood and DNA from samples remaining after the genetic analysis is stored frozen at the biobank of the Biology Department, Uzhhorod National Unive sity, Ukraine. As a result, blood samples were collected from a total 113 individuals.

### b) DNA extraction

Immediately upon arrival to the laboratory, DNA isolation from 200 uL of blood was attempted with innuPREP DNA Blood Minikit (Analitik Gena, Germany). High molecular weight genomic DNA was lightly fragmented by vortexing. The initial DNA concentration was measured with the Implen C40 Nanophotometer (München, Germany), and quality was verified visually on a 2% agarose gel. The 97 successfully extracted DNA samples were normalized to 20-30 ng/μl concentration for downstream application. After the extraction the samples were re-coded and sent to NIH for genotyping procedure, from where the aliquots were further shipped to BGI facility (BGI Shenzhen, CHINA) or to Psomagen Inc. (Gaithersburg, MD, USA) for the whole genome sequencing (WGS). The remaining ∼2 ml was frozen for future use.

### c) Sequencing and Genotyping

All the 97 individuals in this study were sequenced with DNBSEQ-G50 and 88 individuals were cross validated by genotyping using Illumina Global Screening Array. The record of which individual samples have been cross-validated by both technologies is presented in **Table S2**. In addition, a single sample (EG600036) was also sequenced on Illumina HiSeq (∼60x coverage).

#### Sequencing with BGI DNBSEQ-G50

All 97 DNA samples were sequenced on DNBSEQ-G50 (BGI Shenzhen, CHINA). Upon the receipt at the BGI facility, and prior to sequencing, samples were checked again for quality. Concentration was once more detected by fluorometer or Microplate Reader (e.g. Qubit Fluorometer, Invitrogen). Sample integrity and purity were detected by Agarose Gel Electrophoresis (Concentration of Agarose Gel: 1% Voltage:150 V, Electrophoresis Time: 40 min). 1μg genomic DNA was aliquoted and fragmented by Covaris. The fragmented genomic DNA was selected by Agencourt AMPure XP-Medium kit to an average size of 200-400bp. Fragments were end repaired and then 3’ adenylated. Adaptors were ligated to the ends of these 3’ adenylated fragments. PCR products were purified by the Agencourt AMPure XP-Medium kit. The double stranded PCR products were heat denatured and circularized by the splint oligo sequence. The single strand circle DNA (ssCir DNA) was formatted as the final library. The qualified libraries were sequenced by DNBSEQ-G50: ssCir DNA molecule formed a DNA nanoball (DNB) containing more than 300 copies through a rolling-cycle replication. The DNBs were loaded into the patterned nanoarray by using high density DNA nanochip technology. Finally, pair-end 100 bp reads were obtained by combinatorial Probe-Anchor Synthesis (cPAS). Raw reads were filtered removing adaptor sequences, contamination and low-quality reads. Sequencing of all the 97 full genome samples submitted for sequencing at BGI was successful.

#### Short Read Sequencing with Illumina NovaSeek6000

one individual was resequenced by Illumina NovaSeq6000 S4 at Psomagen Inc. (Gaithersburg, MD, USA). Library was prepared using TruSeq DNA PCR Free 350bp protocol by Illumina. The library was sequenced at approximately 64X depth, producing 150bp-long reads, resulting in 241.7G bp of data.

#### Genotyping with the Illumina Infinium Global Screening Array

We attempted to genotype all 97 of the collected samples using the Illumina Infinium Global Screening BeadChip Array-24 v1.0 (GSAMD-24v1-0) for 700,078 loci at the NCI’s DCEG (Bethesda, MD; *https://grcf.jhmi.edu/wp-content/uploads/2017/12/infinium-commercial-gsa-data-sheet-370-2016-016.pdf*). Data was analyzed by using the standard Illumina microarray data analysis workflow. During QC, samples were filtered for contamination, completion rate, and relatedness. As part of QC, we performed ancestry assessment using SNPweights software [44] with a reference panel consisting of 3 populations (European, West African, and East Asian). All samples were attributed to the European ancestry group. After OC and sample exclusion, 87 (86 samples and 1 QC) samples with 689,918 loci and completion rate of 99.9 were retained for further analysis.

### d) Variant Calling

#### Variant Calling of the BGISeq500 data

The sequencing data produced using the DNBSEQ platform for 97 samples were analyzed using the Sention tools (Sentieon Inc, San Jose, CA, USA) high-performance implementation of the BWA/GATK best practices pipeline on servers hosted by the Cornell University Biotechnology Resource Center. Reads were aligned to the GRCh38 human reference genome using BWA-MEM (Version: 0.7.16a-r1181), and mapped reads were prepared for variant calling using Genome Analysis Toolkit (GATK) v3.8-1-0-gf15c1c3ef by Broad), including marking duplicates (*picard MarkDuplicates*, Version 2.12.1), indel realignment (*G*ATK *RealignerTargetCreator, IndelRealigner, Version 3.7-0*), and base quality score recalibration (*GATK BaseRecalib*rator, *PrintReads, Version 3.7-0*). SNP and Indel discovery were performed for each individual using GATK HaplotypeCaller, and merged into a single pVCF using *bcftools*. Sample EG600036 was also run without joint calling which was used when calculating concordance between the Illumina and BGISeq variant callsets. estimated with *TiTvtools* and visualized by *plotTiTv* [18].

#### Repetitive variant calling

Mobile element discovery was performed using MELT (Version 2.2.0) [49] and structural variant discovery using *lumpy-sv* with *Smoove* (Version: 0.2.5)[16]. Short tandem repeats were called using *GangSTR* (Version: 2.4.2) [50] and nuclear mitochondrial DNA using *dinumt [51]*.

### e) Data validation and quality control

Variant files were compared for consistency across the three different platforms: BGI DNBSEQ-G50 sequencing, Illumina genotyping, and Illumina ovaSeq6000 sequencing. Illumina genotyping was performed on 86 of the 97 samples previously sequenced with DNBSEQ-G50. Additionally, one sample (EG600036) was also sequenced with Illumina NovaSeq6000 S4. The variant detection programs were re-run without oint calling for the DNBSEQ-G50 sequencing for sample EG600036 for comparison with the single Illumina sequenced sample. In this sample, the SNPs derived from the WGS platforms were compared to those identified using the Illumina SNP array both for matching position and matching genotype. Structural variants and mobile element insertions were compared between the WGS platforms in EG600036. Variants were considered the same if they had 95% reciprocal overlap. Overall, we found Illumina identified a higher number of larger variants than DNBSEQ-G50. This could potentially be due to its higher coverage (∼60X) compared to DNBSEQ-G50 (∼30X). However, as both have high coverage, we may see diminishing returns for coverage over 30X. An alternative explanation is that the variant identification tools have been built to detect variation from Illumina sequencing data and therefore, may not be able detect variants DNBSEQ-G50 as accurately.

### f) Annotation

Sequence variant files were annotated using *ANNOVAR* [52] and *SNPEff* [53] software using GRCh38 reference databases. The following databases were used for the For ANNOVAR annotations: RefSeq Gene, 1000 genomes superpopulation, dbSNP150 with allelic splitting and left-normalization. For annotation of the medically related and functional variants we used ClinVar version 20200316 [29], InterVar gnomeAd ver 3.0 [11], and *dbnsfp ver. 35c [54]*. For S*NPEff*, the default GRCh38 annotation database [55] was complemented with ClinVar [29] and GWAS catalog [28] database annotation using *snpSift* tool [56].

### g) Population analysis

#### Principal Component analysis (PCA)

For principal component analysis, we used WGS variants of our samples and merged them with samples from neighboring countries available from the European samples from the 1000Genomes Project (Utah Residents (CEPH) with Northern and Western European Ancestry, Toscani in Italy (TSI), Finnish in Finland (FIN), British in England and Scotland (GBR), Iberian Population in Spain (IBS)[13,38]) and French (FRA) and Russians (RUS) from HGDP [39] as well as the relevant high-coverage human genomes Croatian (CRO), Czech (CZ), Estonian (EST), German (GER), Greek (GRE), Hungarian (HUN), Moldovan (MOL), Polish (POL), Russian Cossack (RUS) and Ukrainian (UKR) from the Estonian Biocentre Human Genome Diversity Panel (EGDP) [42], and the Simmons Genome Diversity project [43]. The analysis was performed with *Eigensoft* [47].

To produce a meaningful number of alleles to analyze, the resulting dataset was filtered by genotyping rate (1) and pruned for variants in LD by excluding those with high pairwise correlation within a moving window(--*indep-pairwise* 50 10 0.5). This resulted in 677 samples with 208,945 variants. We used *EIGENSOFT* [47] to calculate the eigenvectors, of which, PC1 and PC2 were visualized using Python programming language, with *pandas, matplotlib and seaborn* libraries [57]. Two extreme outlier samples (EG600056, and EG600052) were left out from the visible range of the PCA plot as they clustered with each other far away from any known European group.

#### Model-based population structure analysis

For the naive (model-based) structure analysis, we used the same dataset described in the Principal Component Analysis (above). The analysis was performed using *ADMIXTURE* software [48]. For identification of the optimal K parameter, we used the 10-fold cross-validation function of *ADMIXTUR*E in range from 2 to 6, with K=3 resulting in the lowest error, deeming it optimal. The results were visualized using Python programming language, with *pandas, matplotlib* and *seaborn* libraries [57,58] to construct a population structure plot using samples from the 1000Genomes Project (Utah Residents (CEU) with Northern and Western European Ancestry, Toscani in Italy (TSI), Finnish in Finland (FIN), British in England and Scotland (GBR), and Iberian Population in Spain (IBS), French population from HGDP(FRA)); [13,38]) and Russians (RUS) from HGDP [39] as well as the relevant high-coverage human genomes from the Estonian Biocentre Human Genome Diversity Panel (EGDP) [42], and Simmons Genome Diversity project [43]. The resulting plot with K=3 is presented in **Figure 3**, and plots with K=4 to K=8 are in the **Figure S3**.

#### Inbreeding estimates

We estimated inbreeding coefficients for all the genotype samples in the same dataset. For this analysis the samples were pruned for genotyping rate (>0.9) and linkage disequilibrium by excluding those with high pairwise correlation within a moving window (plink parameter*--indep-pairwise* 50 10 0.1). Using the resulting dataset containing the remaining 117,641 loci from 84 samples, we performed several inbreeding estimates: (a) method-of-moments F-coefficient estimates, (b) variance-standardized relationship minus 1 estimates, and (c) F-estimates based on correlation between uniting gametes [59]. All the resulting values are presented in **<u>Table S7</u>**, and the estimates for the of method method-of-moments F-coefficient estimates are visualized in a histogram (**Figure S4**).

## Re-use potential

Since the publication of the first human genome [60,61], and the first surveys of worldwide variation such as the 1,000 Genomes project [13,38], the efforts have been directed to expand outwards by expanding the exploration of the human diversity across the world, and filling out more and more “white spots” of genome variation [12,43], as well as inward, to fill the remaining white spots in the human genome itself: to map the remaining gaps in the chromosome assembly and identify new structural and functional variation [62] and to map the three dimensional structure of the human genome [63]. The new data presents a valuable addition to the former and represents the first exploration of the genome landscape in the important component of European genomic diversity.

Genome diversity of Ukraine is an important puzzle to help modern genome studies of population history of Europe. The country is positioned in the crossroad of the early migration of modern humans and the westward expansion of the Indo-Europeans, and represents an aftermath of centuries of migration, admixture, demographic and selective processes. As wave after wave of great human migrations moved across this land for millennia, they were followed by exchange of cultural knowledge and technology along the great trade routes that transect this territory until this day.

The justifications for collecting, sequencing and analyzing populations from this part of Europe has been outlined earlier [10,64], and the new database is a step into that direction. Given its unique history, the genome diversity data from Ukraine will contribute a wealth of new information bringing forth different risk and/or protective alleles that do not exist nor associate with disease, elsewhere in the world. This project identified 13M variants in Ukrainians of which 478 K were novel genomic SNPs currently missing from the global surveys of genomic diversity [11,13]. We also report almost 1M (909,991) complex indels, regions of simultaneous deletions and insertions of DNA fragments of different sizes which lead to net a change in length, with only 713,858 previously reported in gnomAD [11] (**Table 1**). The newly discovered ocal variants can be used to augment the current genotyping arrays and used to screen individuals with genetic disorders in genome wide association studies (GWAS), in clinical trials, and in genome assessment of proliferating cancer cells.

The current project is built upon the open release/access philosophy. The data has been released and can be used to search from population ancestry markers and well as the medically related variants in the subsequent studies. The public nature of the data deposited on the specially created web resource located at Uzhhorod National University, will ensure that the biomedical researchers in the country will receive access to a useful information resource for future projects in genomics, bioinformatics and personalized medicine. Engaging local Ukrainian scientists in this collaborative international project like building the foundation for the future studies and ensuring their participation in the worldwide research community.

## Availability of source code and requirements

### Availability of the Supporting Data

The raw reads are available at the SRA (Project PRJNA661978, SUB7904361). All other databases mentioned in this project are available in GigaDB.

## List of Supplementary Tables (available in GigaDB)

**<u>Table S1</u>**. Sequencing summaries of output from DNBSEQ-G50 and Illumina NovaSeq6000. Full sequencing statistics for individual samples in **<u>Table S1.2</u>**

**<u>Table S2.</u>** Filtering summary of the data obtained from 97 whole genomes sequenced with DNBSeq-G50

**<u>Table S3</u>**. The full list of high impact functional variants (including frameshift, start lost/stop lost or gained, transcript ablations and splice alterations) that had an allele count of two or more with their predicted function, number of gene transcripts of the gene affected, and frequencies.

**Table S4**. List of the medically relevant functional markers found in the Ukrainian population and reported in A. **<u>GWAS catalog</u>** [28] and B. **<u>ClinVar</u>** [29] databases. Allele frequency is reported compared to the reference allele in GRCh38.

**<u>Table S5</u>**. Complete list of the highly differentiating markers, reported in ClinVar [29], with high differences in the Ukrainian population compared to the neighboring populations in other European populations (the combined sample from Western and Central Europe from 1000Genomes Project with French samples from HGDP (EUR)[13,38,39] and Russians (RUS) from HGDP [39]. Non-reference allele frequency (NAF) is reported compared to the reference allele in GRCh38. Differences are evaluated by the Fisher Exact Test (FET).

**<u>Table S6</u>**. A list of markers with the highest non-reference allele frequency (NAF) differences in the Ukrainian population evaluated by the Fisher Exact Test (FET) compared to the frequencies in the neighboring populations: the combined population from Europe (EUR) [13] and Russians (RUS) from HGDP [39]. This database contains candidate ancestry informative markers (or AIMs)[44], that can be used for mapping disease alleles by admixture disequilibrium [45,46].

**<u>Table S7</u>**. Inbreeding estimates in a dataset of 117,641 loci from 84 samples: (a) method-of-moments F-coefficient estimates, (b) variance-standardized relationship minus 1 estimates, and (c) F-estimates based on correlation between uniting gametes [59].

## List of Supplementary Files (available in GigaDB)

**Supplementary File 1**. IRB approval of the study “Genomic Diversity of Ukraine’s Population” (*in Ukrainian*). <u>Supplementary File 1. The IRB Approval.jpg</u>

**Supplementary File 2**. Genomic Diversity of Ukraine’s Population Project: Protocol description, questionnaire, and informed consent to participate and publish (*in Ukrainian with English Transla*tion). <u>Supplementary File 2. The Informed Consent</u>

**Supplementary File 3**. The list of the samples in this study, their characteristics and geographical locations, and sources of genomic data for each (DNBSEQ-G50 sequencing (BGI Inc., Shenzhen, China), Illumina Global Screening Array genotyping, and Illumina HiSeq sequencing array (Illumina Inc., San Diego, USA). <u>Supplementary File 3. The List of Samples</u>

**Supplementary File 4**. The full annotation database with classifications of variants in the Ukrainian populations from 97 genomes fully sequenced on BGISeq500. *ftp://user81@8.210.79.81/Ukraine_bgi_all_ann_GWAS_Clinvar.vcf.gz*

**Supplementary File 5**. List of the samples from different studies used in the current population analysis. <u>Supplementary File 5. Sample Sources</u>

## Supplementary Figures

**Figure S1.**
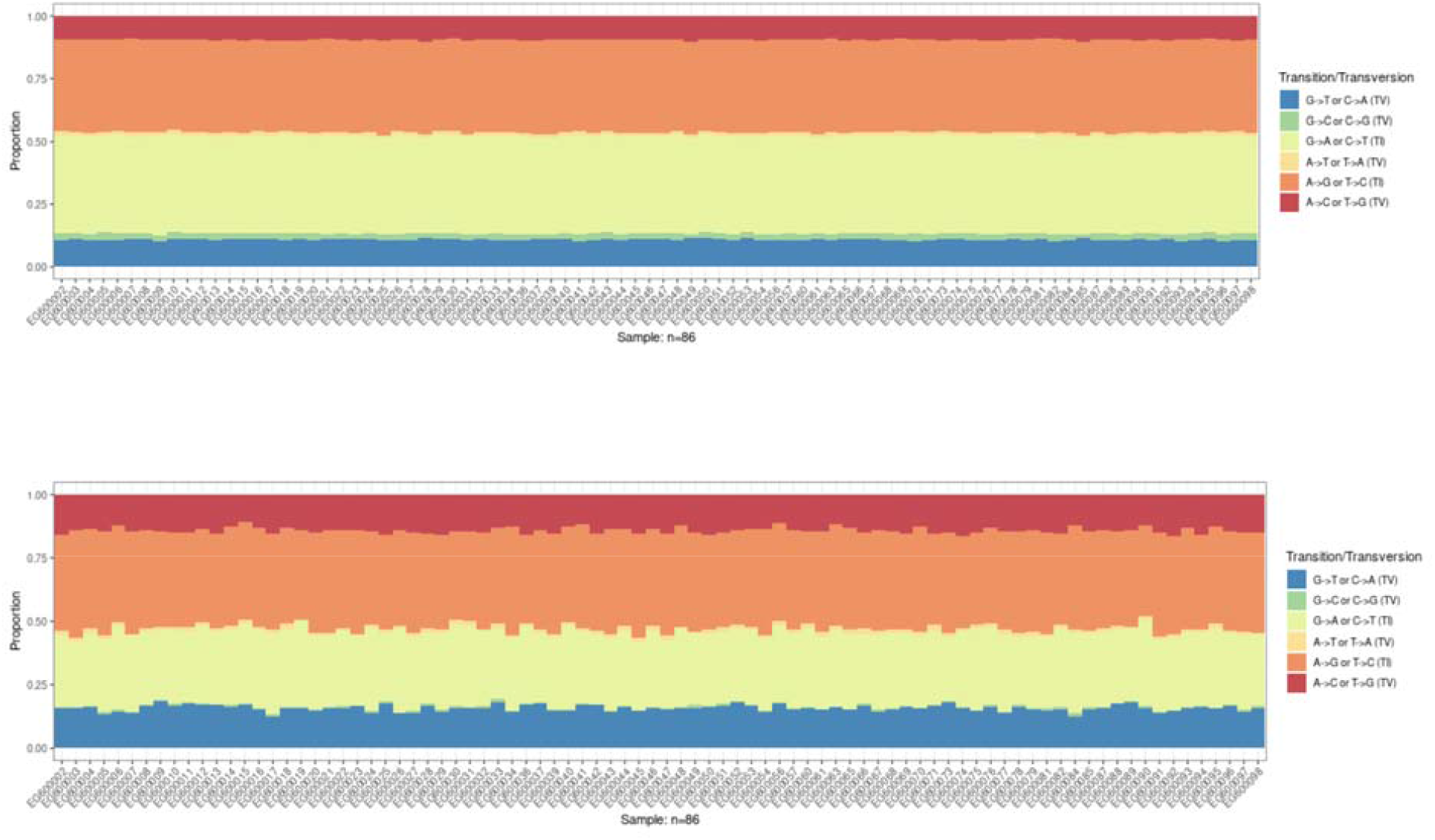
Transition/Transversion ratio (or TITV ratio) for the novel SNPs (estimated with *TiTvtools* [18] and visualized by *plotTiTv*) (top) for the SNPs where Illumina SNP array identified more alternate haplotypes than BGI (top right triangle in Figure 1C) and (bottom) for the SNPs where BGISeq identified more alternate haplotypes than Illumina SNP Array (bottom left triangle on Figure 1C table).

**Figure S2.**
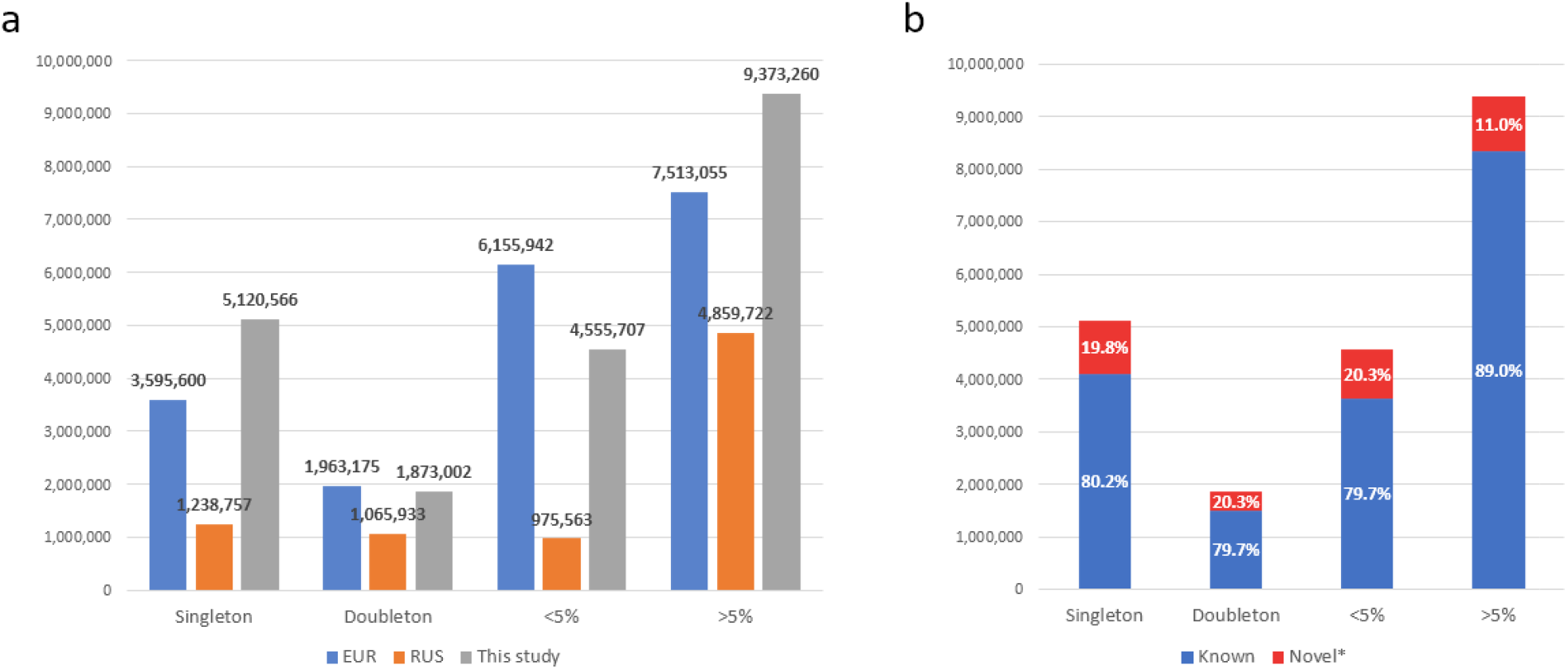
**A.** Frequencies of various classes of SNPs in the Ukrainian genome variation database. Definitions are as follows: Singleton (passed the GATK QC once), Doubleton, Rare (3-10 counts roughly equivalent to 1%< x < 5%) and Common (>5%) to make it closer to the 1KGP definitions. B. Percent novel mutations in various classes of SNPs.

**Figure S3.**
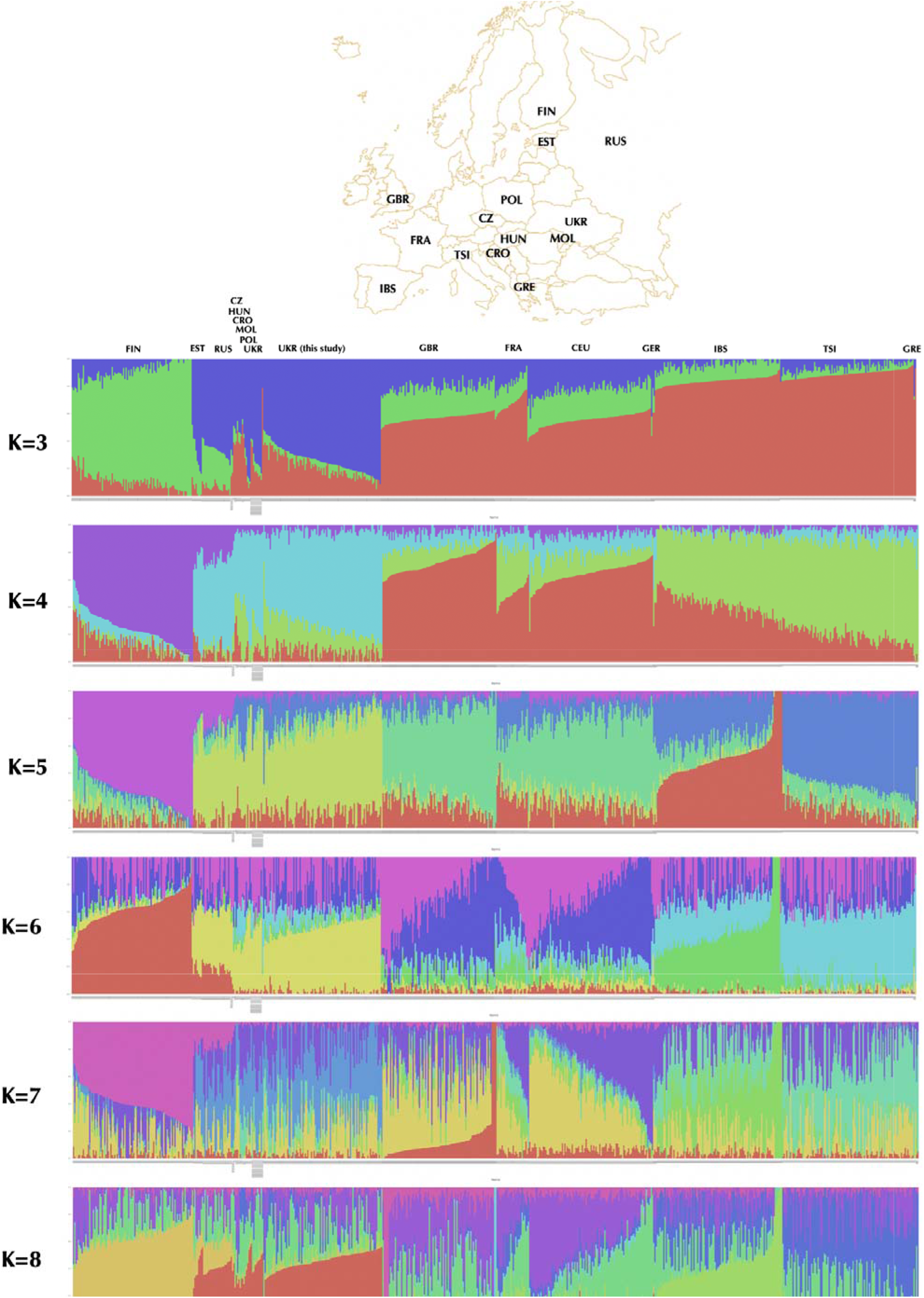
Genetic structure of Ukrainian population in comparison to other European populations. For identification of the optimal K parameter, we used the 10-fold cross-validation function of *ADMIXTURE* in range from 2 to 8, with K=3 resulting in the lowest error [48]. This analysis included genomes from this study as well as samples from the 1000Genomes Project (Utah Residents (CEU) with Northern and Western European Ancestry, Toscani in Italy (TSI), Finnish in Finland (FIN), British in England and Scotland (GBR), and Iberian Population in Spain (IBS)[13,38], French(FRA) and Russians (RUS) from HGDP [39], as well as the relevant high-coverage human genomes Croatian (CRO), Czech (CZ), Estonian (EST), German (GER), Greek (GRE), Hungarian (HUN), Moldovan (MOL), Polish (POL), Russian Cossack (RUS) and Ukrainian (UKR) from the Estonian Biocentre Human Genome Diversity Panel (EGDP) [42] as well as Simmons Genome Diversity project [43].

**Figure S4.**
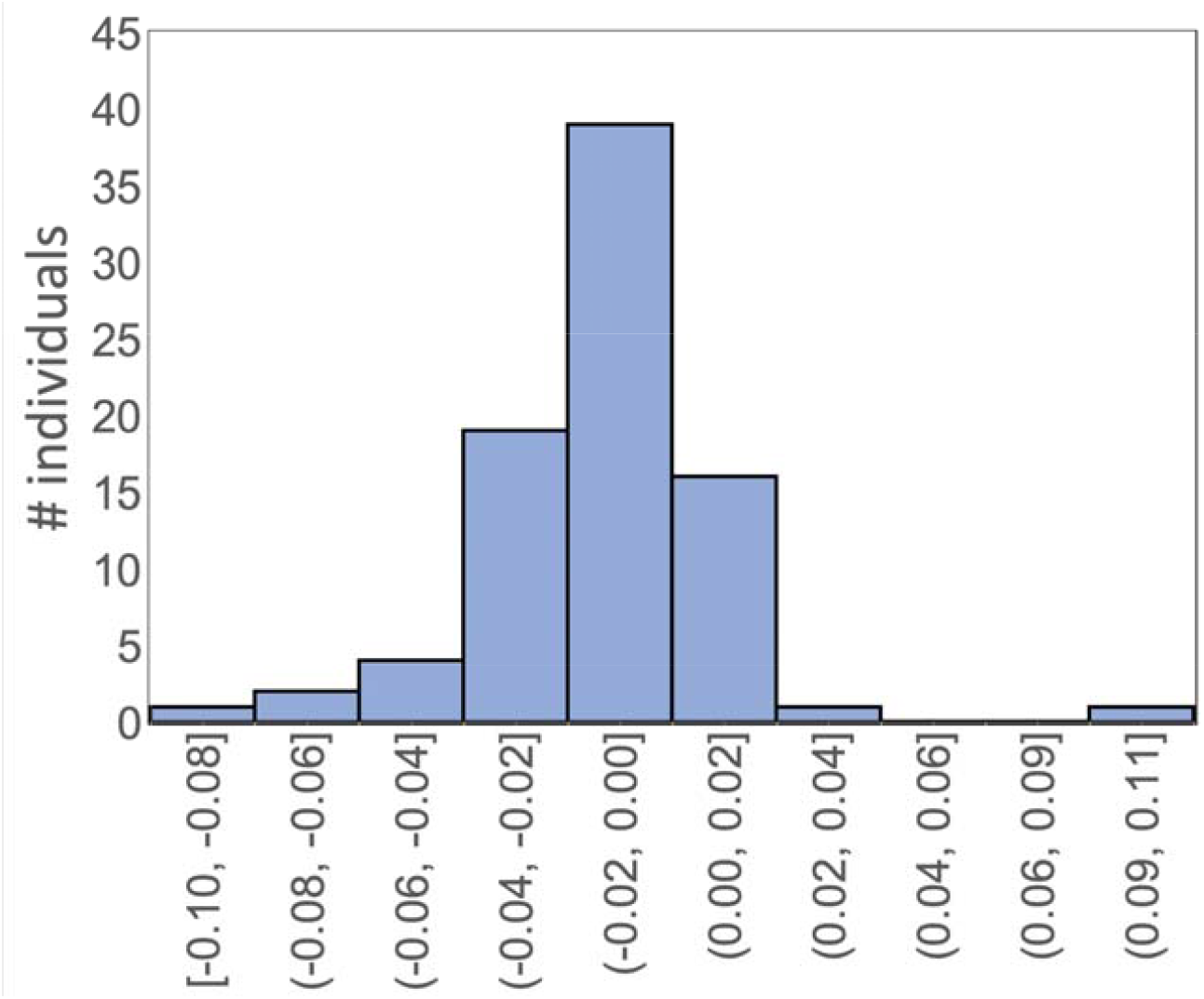
Distribution of inbreeding coefficients in the Ukrainian sample. The individual values corresponding to the samples are presented in **Table S7**

## Supplementary Tables

**Table S1.** Sequencing summary of output from DNBSEQ-G50 and Illumina NovaSeq6000.

|  | **DNBSeq-G50** <sup>□</sup> | **Illumina NovaSeq6000** <sup>¥</sup> |
| --- | --- | --- |
| **Samples sequenced** | 97 | 1 |
| **Read length (bp)** | 100 | 150 |
| **Reads above Q20**<br>**(>99% quality score)** | 97.85% | 96.91 % |
| **Total Reads** | 99,638,538,182 | 1,600,898,738 |
| **Average reads/sample** | 1,027,201,425 | 1,600,898,738 |
| **Average GC content** | 42.05% | 41.07 |

<sup>□</sup> Sequencing of 97 samples were attempted on DNBSeq-G50 at BGI sequencing facility (BGI Shenzhen, CHINA), and all 97 were successful.
<sup>¥</sup> One sample (EG600036) was sent to Illumina NovaSeq6000 S4 at Psomagen Inc. (Gaithersburg, MD, USA). In addition, 96 samples were genotyped using Illumina Global Screening Array array (Illumina Inc., San Diego, USA), and 87 were successful (86 individual samples and 1 internal QC) remained after filtering.

**Table S2.** Filtering summary of the data obtained from 97 whole genomes sequenced with DNBSeq-G50.

| Sequencing results | All samples |  |  |
| --- | --- | --- | --- |
|  | Total Unique SNPs # | Filtered Count | % Filtered <sup>□</sup> |
| **Variation** |  |  |  |
| **SNPs** | 14,738,063 | 1,727,084 | 11.7 |
| **Bi-allelic** | 14,254,070 | 1,586,787 | 11.1 |
| **Multi-allelic** | 483,993 | 140,297 | 29.0 |
| **Small Indels <sup>¥</sup>** | 2,808,384 | 80,780 | 2.9 |
| **Deletions** | 1,864,698 | 57,959 | 3.1 |
| **Insertions** | 1,488,408 | 42,421 | 2.9 |
| **Structural Variants <sup>\$</sup>** | | | |
| **Large Deletions** | 685,56 | 52,478 | 76.5 |
| **Large Duplications** | 3,374 | 52,478 | 45.3 |
| **Inversions** | 430 | 93 | 21.6 |
| **Mobile Element Insertions** |  |  |  |
| **Alu** | 7550 | 1790 | 23.7 |
| **L1** | 3123 | 2672 | 85.6 |
| **SVA** | 222 | 122 | 55.0 |
| **NUMT** | 1169 | 455 | 38.9 |

## Declarations

### List of abbreviations

#### Consent for publication

The collection procedure was approved as part of the “*Genome Diversity in Ukraine”* project by the Institutional Review Board (IRB) of Uzhhorod National University, Uzhhorod Ukraine (**Supplementary File 1**). Each participant had an opportunity to review the informed consent (**Supplementary File 2**), have been explained the nature of the genome data and take a decision about making it public.

#### Competing interests

The following authors declare that they have no competing interests:

Taras K. Oleksyk, Walter W. Wolfsberger, Alexandra Weber, Khrystyna Shchubelka, Stephanie O. Castro-Marquez, Sarah Medley, Alina Urbanovych, Patricia Boldyzhar, Viktoriya Stakhovska, Kateryna Malyar, Yaroslava Hasynets, Juan L. Rodriguez-Flores, Fabia Battistuzzi, Siru Chen, Meredith Yeager, Michael Dean, Olga T. Oleksyk, Ryan E. Mills, and Volodymyr Smolanka

The following authors may have competing interests:

Yuan Liu, Huanming Yang, Olga Levchuk, Alla Patrus, Nelya Lazar

#### Funding

This research was funded in part by the internal funding from BGI (China), Uzhhorod National University (Ukraine), Division of Cancer Epidemiology and Genetics, National Cancer Institute (USA), and the startup fund of Oakland University, Rochester, Michigan.

### Authors’ contributions

**Conceptualization:** Taras K. Oleksyk, Khrystyna Shchubelka, Walter Wolfsberger, Siru Chen, Huanming Yang, Yuan Liu, Volodymyr Smolanka, Juan L. Rodriguez-Flores, Fabia Battistuzzi, Olga T. Oleksyk, Michael Dean, Meredith Yeager, Ryan Mills, and Volodymyr Smolanka

**Data curation:** Walter Wolfsberger, Khrystyna Shchubelka, Alexandra Weber, Alina Urbanovych, Patricia Boldyzhar, Viktoriya Stakhovska, Kateryna Malyar, Yaroslava Hasynets, Nelya Lazar, Olga T. Oleksyk

**Formal analysis:** Walter Wolfsberger, Alexandra Weber

**Funding acquisition**: Taras K. Oleksyk;

**Investigation:** Khrystyna Shchubelka, Walter Wolfsberger, Alexandra Weber, Stephanie Castro-Marquez;

**Methodology:** Taras K. Oleksyk, Michael Dean, and Ryan Mills

**Project administration**: Taras K. Oleksyk, Michael Dean, and Volodymyr Smolanka;

**Resources:** Taras K. Oleksyk, Huanming Yang, Yuan Liu, Ryan Mills, Meredith Yeager, Michel Dean, Olga T. Oleksyk, and Volodymyr Smolanka

**Software:** Walter Wolfsberger, Alexandra Weber;

**Supervision:** Taras K. Oleksyk, Volodymyr Smolanka, Michael Dean, and Ryan Mills;

**Visualization:** Walter Wolfsberger, Khrystyna Shchubelka, Alexandra Weber;

**Writing**, Taras K. Oleksyk, Khrystyna Shchubelka, Walter Wolfsberger, Alexandra Weber (original draft), and Taras K. Oleksyk and Ryan Mills (review & editing).

#### Acknowledgements

We thank all the Ukrainian volunteers who contributed their data for the project.

## Group authorship

n/a

## Authors’ information

n/a

## Endnotes

n/a

## Web links and URLs

*GigaDB*

*http://genomes.uzhhorod.edu.ua*

